# *Brucella* outer membrane proteins to control maturation and antigen presentation of mouse dendritic cells

**DOI:** 10.1101/2021.03.05.434055

**Authors:** Ningning Yang, Zhixia Tong, Zhen Wang, Mingguo Xu, Qian Zhang, Huan Zhang, Yueli Wang, Jihai Yi, Tianhao Sun, Buyun Cui, Chuangfu Chen

## Abstract

Brucellosis is a highly contagious zoonotic disease, which seriously endangers animal husbandry in China. Bone marrow-derived dendritic cells (DCs) are full-time antigen presenting cells (APC) that play an important role in the interaction between pathogens and host immunity. DCs were stimulated with *Brucella* major outer membrane proteins (OMPs: OMP10, OMP19, OMP25, BP26 and OMP31) and *Brucella* mutants (Δ *omp10*, Δ *omp19*, Δ *omp25*, Δ *bp26*, Δ *omp31*) to examine effects on DC maturity and antigen presentation. *Brucella* OMP10, OMP19 and BP26; *Brucella* mutants Δ *omp10*, Δ *omp19*, Δ *omp25*, Δ *bp26*, Δ *omp31* and *Brucella* RB51 induced DC maturation and antigen presentation efficiency in mice, activated proliferation of T lymphocytes, and decreased apoptosis, which helped the host recognize antigens and eliminate pathogens. However, *B. abortus* 2308 evaded the host immune function and established chronic infection by maintaining a balance between intracellular replication and inducing apoptosis, thus reducing DC maturation and antigen presentation to T cells. Toll-like receptor (TLR) -mediated signaling pathways were involved in the DC maturation and antigen presentation induced by *Brucella* OMPs. These results enhance understanding of *Brucella* pathogenesis and the host protective immune response mechanism and lay the foundation for the rational design of *Brucella* vaccines.

## Introduction

Brucellosis is one of the most common zoonotic diseases in the world. This chronic infectious disease can affect the reproductive function of animals and may lead to abortion and infertility, resulting in huge economic losses [1]. Current studies have shown that brucellosis generally does not spread from person to person, however because it can be transmitted via aerosol, there is a high global incidence. Thus, the World Organization for Animal Health (OIE) classifies brucellosis as a class B epidemic disease [2-4] with more than 500,000 people infected each year [5,6].

The main outer membrane component of *Brucella* is composed of lipopolysaccharide (LPS) and outer membrane proteins (OMPs). The OMP of *Brucella* is described as a “complex structure filled by at least 75 proteins” [7,8]. In the past 10 years, several kinds of OMPs have been characterized on a molecular level and some studies have been done, however, the research has not been sufficiently depth. *Brucella* OMPs are the initial contact point between the pathogen and the host immune system and may potentially protect against *Brucella* infection by inducing cell-mediated immunity [9,10]. DCs are considered to be among the most effective antigen-presenting cells in the immune system. They are the only cells that can activate the initial T lymphocytes, thus triggering the adaptive immune response of the body and are the most effective professional antigen-presenting cell connecting innate immunity and acquired immunity [11-13]. Considering that DCs play an irreplaceable role in the initiation and orientation of the acquired immune response, establishing the model of interaction between *Brucella* and DCs may be more suitable than the traditional macrophage model for understanding the relationship between the pathogenic mechanism of *Brucella* and the specific immune response of the infected host. However, several pathogens, including *Brucella*, have evolved to maintain DCs in a semi-mature state to escape recognition by the immune system. In vivo studies have shown that *Brucella* can effectively proliferate in DCs, and in vitro studies have demonstrated that *B. abortus* 2308 and *B. suis* 1330 can inhibit the phenotype maturation of DCs [14,15]. In fact, *Brucella* infection in DCs is characterized by a weak expression of major histocompatibility complex class II (MHC-II) and the costimulatory molecules CD80 and CD86 on the surface of DCs, leading to the inhibition of the secretion of inflammatory cytokines (TNF-α, IL-12), ultimately hampering antigen presentation by DCs to T lymphocytes.

The purpose of this study was to investigate the effects of *Brucella* OMPs on the DC maturity and antigen presentation, as well as on innate and acquired immunity, so as to lay a theoretical foundation for the rational design of a brucellosis vaccine and to further understand the pathogenic mechanism of brucellosis and the protective immune response mechanism of the host.

## Materials and Methods

### Animals and Feeding

All the experimental procedures involving animals were approved by the Animal Experimental Ethical Committee Form of the First Affiliated Hospital of Medical College, Shihezi University. Ethical Committee Approval Notice No. A 280-167. Female BABL/C mice, 6–8 weeks old with mean BW of 20 ± 5 g were purchased from the Animal Experiment Center of Xinjiang Medical University. All of mice were given enough food and water, a 12 h light-dark cycle and 15–20°C, 50% relative humidity.

### *Bacteria* strains

*B. abortus* 2308, *B. abortus* RB51 were obtained from the Center of Chinese Disease Prevention and Control (Beijing CDC, China), *Brucella* PET-30a-omp10[16], PET-32a-omp19[17], PET-28a-omp25[18], PET-28a-bp26[19], PET-30a-omp31[20] the prokaryotic recombinant expression strain and *Brucella* mutants Δ*omp10*[21], Δ*omp19*[22], Δ*omp25*[23], Δ*bp26*[24] and Δ*omp31*[25] were all constructed by the zoonotic laboratory of Shihezi University. Each bacterial strain was cultured in tryptone soya agar (TSA) (Becton, Dickinson and Company, USA).

### Acquisition of OMPs

OMP10, OMP19, BP26, OMP25, OMP31 were obtained by His-Tagged Protein Purification Kit (Inclusion Body Protein) (Kangwei, China) (Fig. S1). All the proteins were endotoxin free (ToxinEraserTM Endotoxin Removal Kit, GenScript Biotechnology, Nanjing, China).

### Verification of Brucella mutants

*Brucella* mutants (Δ*omp10*, Δ*omp19*, Δ*omp25*, Δ*bp26* and Δ*omp31*) were indentified by PCR with corresponding identification primers (Table S1 and Fig. S2)

### Isolation and cultivation of DCs

The mouse DC isolation method of Inaba [26-28] et al was used in this study. Briefly, cells were flushed and purified from the tibias and femurs of BABL/C mice, which had been euthanized by cervical dislocation. The cells were plated at 1×10^6^/well in 2 mL of RPMI 1640 and then cultured at 37°C in 5% CO_2_ for 3 h. The supernatant was then discarded to remove non-adherent cells, and fresh 10% fetal bovine serum (10% FBS) with GM-CSF (10 ng/mL) and IL-4 (10 ng/mL) (all PeproTech, USA) were added. Fresh medium was added on days 2 and 4 and cells were collected on day 6 (Fig S3). Flow cytometry was used to identify PE-anti-mouse CD11c, PE-anti-mouse CD80, PE-anti-mouse CD83, PE-anti-mouse CD86, PE-anti-mouse CD40, PE-anti-mouse MHC-I and PE-anti-mouse MHC-II (1 μL/tube) (BioLegend, California, USA) (Fig. S4).

### Flow cytometry analysis

DCs treated with OMPs and *Brucella* mutants were analyzed via flow cytometry for expression of multiple markers of maturation on the cells surface. The following antibodies and appropriate isotype controls (all BD Biosciences, USA) were utilized. DCs were stimulated by purified OMPs (50 ug/mL) (endotoxin removed), *Brucella* LPS groups (R-LPS, S-LPS) (1 ug/mL), Escherichia coli (*E*.*coli*) LPS (E-LPS) (1 ug/mL), or *Brucella* mutants at a multiplicity of infection (MOI) of 5, PBS was used as a control.

After 24 h, the cells were collected and washed three times and the cell concentration was adjusted to 1 × 10^6^ cells/tube using PBS. All antibodies (PE-anti-mouse CD11c, PE-anti-mouse CD80, PE-anti-mouse CD83, PE-anti-mouse CD86 and PE-anti-mouse CD40 (1 μL/tube)) were added to each tube. The DCs and antibodies were mixed and incubated at 4°C for 20 minutes and then washed three times and the unbound antibodies were removed. The cells were resuspended in PBS (300 mL) and cell phenotype changes were detected by flow cytometry [14,29]. Data were collected on a FACSCalibur (BD Biosciences) and analyzed using FlowJo software (Tree Star).

### Cytokine detection

Cytokine measurements [9,30] were done at 24 h. The cell culture supernatant was collected and the expression of cytokines TNF-α, IFN-γ, IL-6, IL-12, IL-10 and IL-4 were quantified according to ELISA kit instructions (BD Biosciences).

### Quantitative real-time PCR

Expression of transcriptional factors including TLR2, TLR4 and TLR9 mRNA were evaluated using real-time PCR (RT-PCR). Briefly, the total RNA (1 µg) was extracted using TRIzol reagent and reverse transcribed to synthesize cDNA (Kangwei, China). The standard curve was drawn for the genes from cDNA. GADPH was used as the endogenous control and specific primers for this gene were designed using Primer 5 software (Table 1). Comparative RT-PCR (Takara, Japan) was performed using SYBR Green Supermix via an RT-PCR machine (RocheLightCycler480, Germany). Finally, TLR2, TLR4 and TLR9 relative expressions were determined using the equation 2-ΔCT [31,32].

**Table 1.**
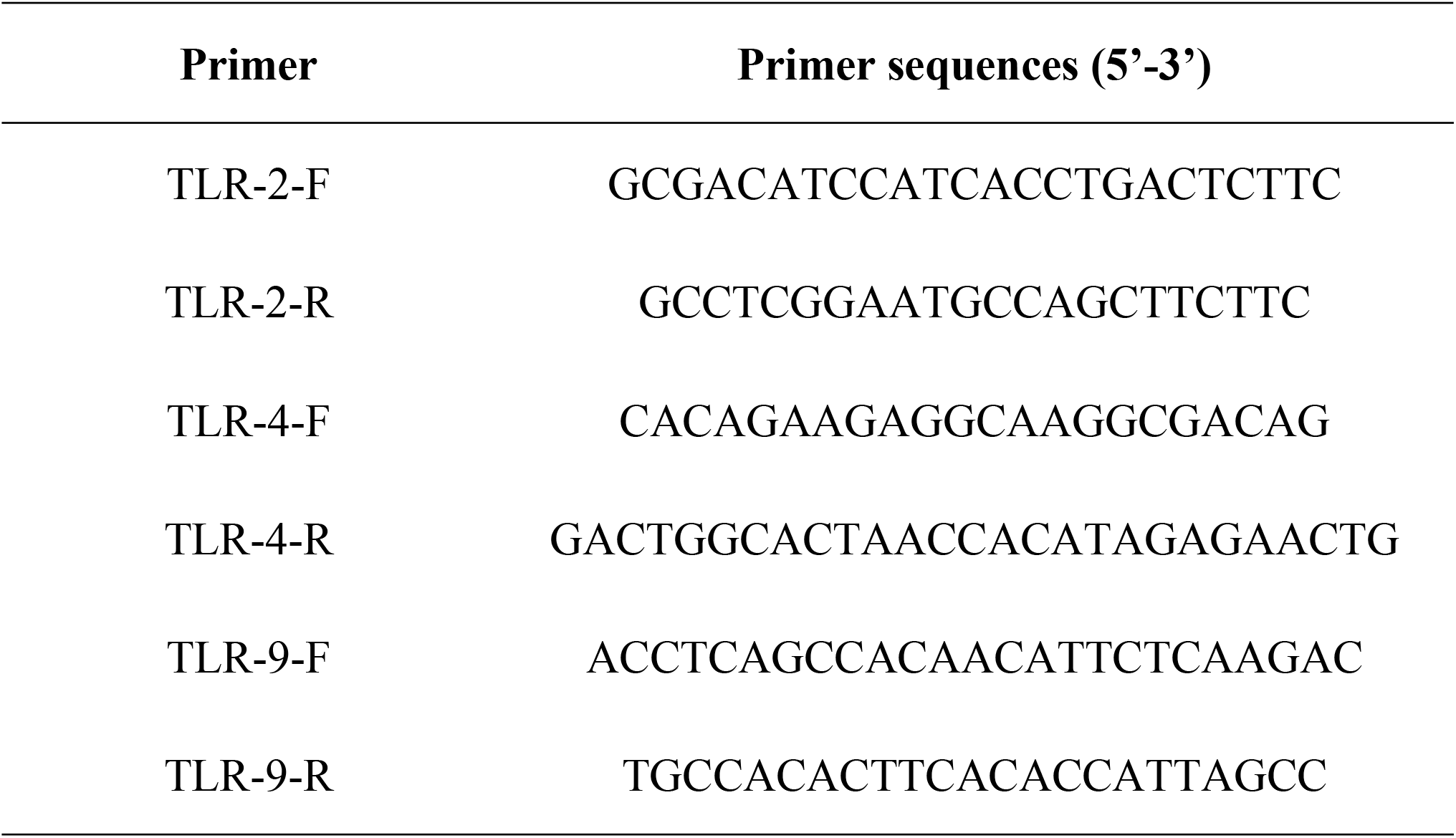
Primers and sequences of TLRs

### Cell viability assay

DCs (day 6) were collected and the cells were stimulated with OMPs and *Brucella* mutants. After 48 h, the stimulated cells were collected and the concentration was adjusted to 1 × 10^5^ cells/mL in RPMI-1640 medium. The negative and positive control groups were given the same treatments. The cells were mixed with mitomycin C (25 μg/mL) and incubated for 30 min at 37 °C. Cells were then placed into a 96-well cell culture plate (100 μL per well) as stimulating cells. Mouse spleen lymphocytes were prepared using a Solebo mouse spleen lymphocyte separation kit (Solebo, Beijing, China). The cells were obtained, the concentration was adjusted to 5.0 × 10^6^ cells/well, and 5.0 × 10^4^ cells were added to each well of the 96-well cell culture plate as reactive cells. The ratios of stimulated cells to reactive cells were 1: 25, 1: 50 and 1: 100, with three wells for each ratio. After 72 h, MTT (20 μL) was added to each well, and the cells were incubated for 4 h. The intracellular punctate particles were observed under microscope. The supernatant was discarded and DMSO (100 μL) was added to each well and the plate was incubated in the dark for about 4 h. After the purple crystal was completely dissolved under the microscope, the OD_570_ nm value was detected. The stimulation index (SI) = OD value of stimulated group/OD value of control group [33-35]. The experiment was repeated four times.

### Apoptosis detection

The Annexin V-FITC apoptosis assay kit (Absin Bioscience Inc, China) was used to evaluate the rate of apoptosis in MCs. DCs (5 × 10^5^ cells) were infected with *Brucella* and *Brucella* mutants at a MOI of 5. After 24 h, DCs were collected and were washed three times with PBS. A 100 μL volume of the binding buffer was added to each tube to resuspend the cells. Annexin V-FITC (5 μL) was added to the binding buffer, the solution was mixed well, and the tubes were incubated for 15 min in the dark at room temperature. Propidium iodide (5 μL) was added and the solution mixed well 5 min prior to flow cytometry. Binding buffer (200 μL) was then added to each tube and the cell apoptosis assay was performed using the flow cytometry apparatus (BD, USA). Each experiment was performed three times [36,37].

### Intracellular survival

DCs were infected with *Brucella* and *Brucella* mutants at a MOI of 5. At 45 min, DCs were washed three times with PBS and then incubated with 50 µg/mL gentamicin (Invitrogen Life Technologies, USA) for 1 h to kill the remaining extracellular bacteria. DCs were subsequently washed three times with PBS and cultured with fresh 10% FBS. DCs were collected after 1, 6, 12, 24 and 48 h of infection. DCs were then washed three times, 5 min each time, with PBS and then cleaved with TritonX-100 (0.1%). After continuous gradient dilution to a suitable concentration, 100 μL lysate was used to coat a TSB Agar *Brucella* plate (three repeated controls for each dilution) and cultured (inverted) at 37°C in a 5% CO_2_ incubator for 3–4 days to determine the number of CFU (CFU/well) [38].

### Statistical analysis

Data are presented as mean ± standard deviation and were analyzed using Graph Pad Prism software (Graph-Pad Software Inc, San Diego, CA, USA). The differences groups were analyzed by the analysis of variance using SPSS 17.0 software (SPSS, Inc. Chicago, IL, USA). A *P*-value of < 0.01 was considered greatly significant, and a *P*-value of < 0.05 was considered significant. *** *P* < 0.001. All experiments were independently performed at least three times.

## Results

DC maturation and antigen presentation were affected by OMPs and *Brucella* mutants Effects of OMPs and *Brucella* mutants on cell maturation were assessed using flow cytometry. PE-anti-mouse CD40, PE-anti-mouse CD80, PE-anti-mouse CD86, PE-anti-mouse CD83, PE-anti-mouse MHC-I and PE-anti-mouse MHC-II (Abbreviations: CD40, CD80, CD86, CD83, MHC-I and MHC-II) were found to be significantly increased in the OMP10, OMP19, BP26 and *Escherichia coli* LPS (E-LPS) (*P* < 0.01), and CD86 and MHC-II were markedly enhanced in *Brucella* LPSs group (R-LPS: RB51-LPS; S-LPS: 2308-LPS) (*P* < 0.05), whereas CD40, CD86, MHC-I and MHC-II were obviously decreased in the OMP25 and OMP31 (*P* < 0.01) (Fig. 1A) compared with the phosphate-buffered saline (PBS) control group. CD40, CD80, CD86, CD83, MHC-I and MHC-II were significantly increased in the *Brucella* mutants (Δ*omp10*, Δ*omp19*, Δ*omp25*, Δ*bp26*, Δ*omp31*) and *B. abortus* RB51 (*P* < 0.05). Finally, the *B. abortus* 2308 significantly inhibited expression of all of these surface markers in DCs (*P* < 0.01) (Fig. 1B). These results demonstrate that *Brucella* OMPs and *Brucella* mutants can affect the maturation and antigen presentation of DCs.

**Fig. 1.**
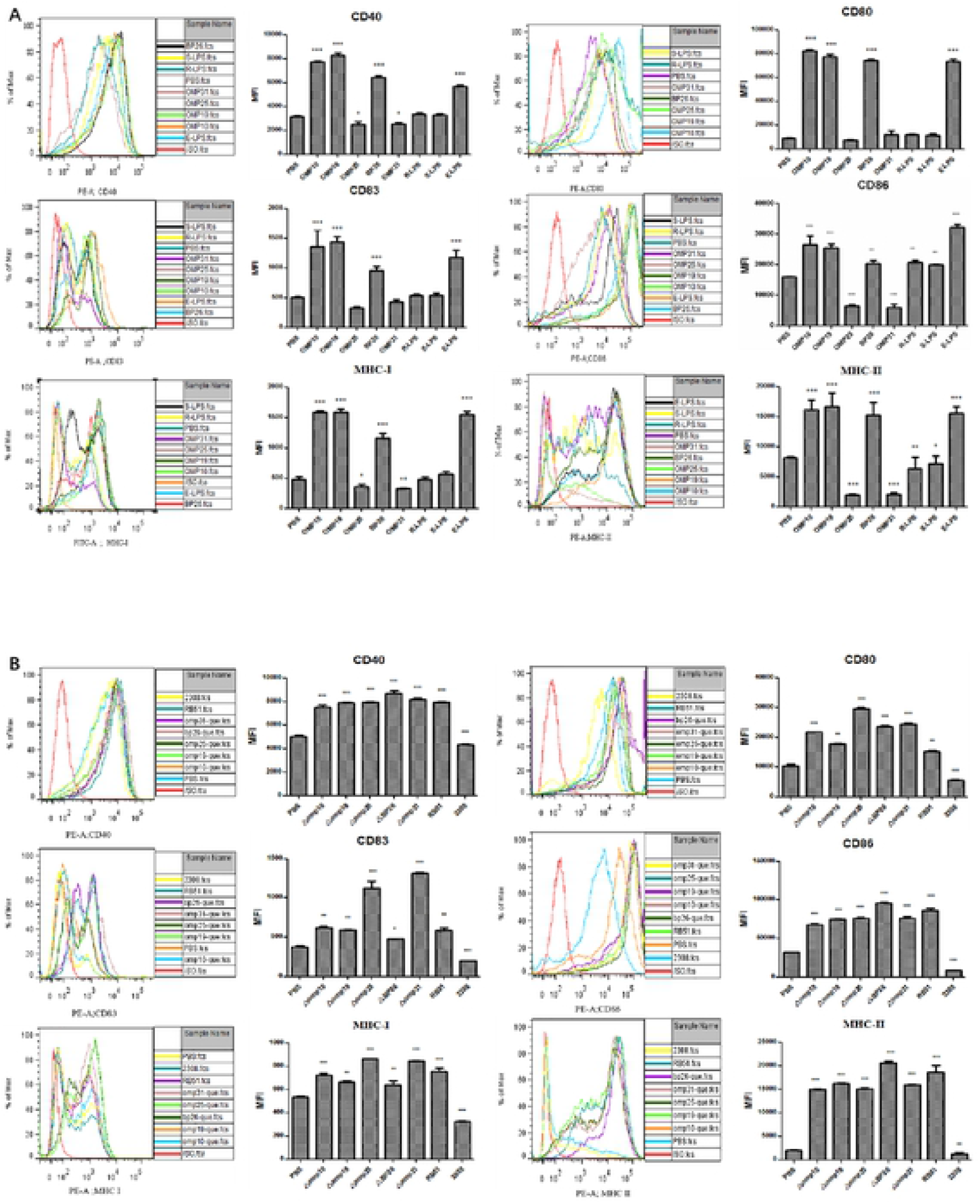
The effects of *Brucella* major OMPs and *Brucella* on surface molecule expression in DCs. DCs were cultured by purified outer membrane proteins (OMPs: OMP10, OMP19, OMP25, BP26, and OMP31) (endotoxin removed), R-LPS, S-LPS, E-LPS(A), or *Brucella* mutants (Δ*omp10*, Δ*omp19*, Δ*omp25*, Δ*bp26*, Δ*omp31*), *B. abortus* RB51, 2308 and PBS for 24 h (B). DCs were harvested and analyzed by flow cytometry. Cells were stained with fluorescent antibodies against CD80, CD83, CD86, CD40, MHC-I and MHC-II. On the left is a histogram of the presented fluorescence as measured by flow cytometry, and on the right is a histogram of the mean fluorescence intensity (MFI) obtained by statistical analysis. The data are representative of three independent experiments. \**P* < 0.05, \*\**P* < 0.01, and \*\*\**P* < 0.001. R-LPS: RB51-LPS; S-LPS: 2308-LPS; Escherichia coli LPS: E-LPS; phosphate-buffered saline: PBS.

### The secretion of cytokines was affected by OMPs and *Brucella* mutants

To analyze the effect of OMPs and *Brucella* mutants on cytokine secretion, the levels of IL-4, IL-6, IL-10, IL-12, TNF-α and INF-γ were measured in the supernatant via ELISA. Compared with the control group, the levels of IL-6, IL-12, INF-γ and TNF-α were dramatically increased in the OMP10, OMP19 and BP26 (*P* < 0.01), whereas the levels of IL-4 and IL-10 were significantly decreased in the OMP10, OMP19, BP26 and LPS (*P* < 0.01). The levels of IL-4 and IL-10 were obviously enhanced in the OMP25 and OMP31 (*P* < 0.05), and the levels of IL-6, IL-12 and TNF-α were reduced (*P*<0.05) (Fig. 2A). The *Brucella* mutants (Δ*omp1*0, Δ*omp19*, Δ*omp25*, Δ*bp26*, Δ*omp31*) and B. abortus RB51 infection of DCs significantly induced the secretion of IL-6, IL-10, IL-12 and TNF-α (*P* < 0.01) and significantly inhibited the secretion of IL-4 and IL-10 (*P* < 0.001). *B. abortus* 2308 infection significantly inhibited the secretion of IL-6, IL-12 and INF-γ (*P* < 0.05), but significantly increased the secretion of IL-4 and IL-10 (*P* < 0.001) (Fig. 2B). These results show that the cytokine secretion was affected by OMPs and *Brucella* mutants, these results are consistent with the DC maturation results.

**Fig. 2.**
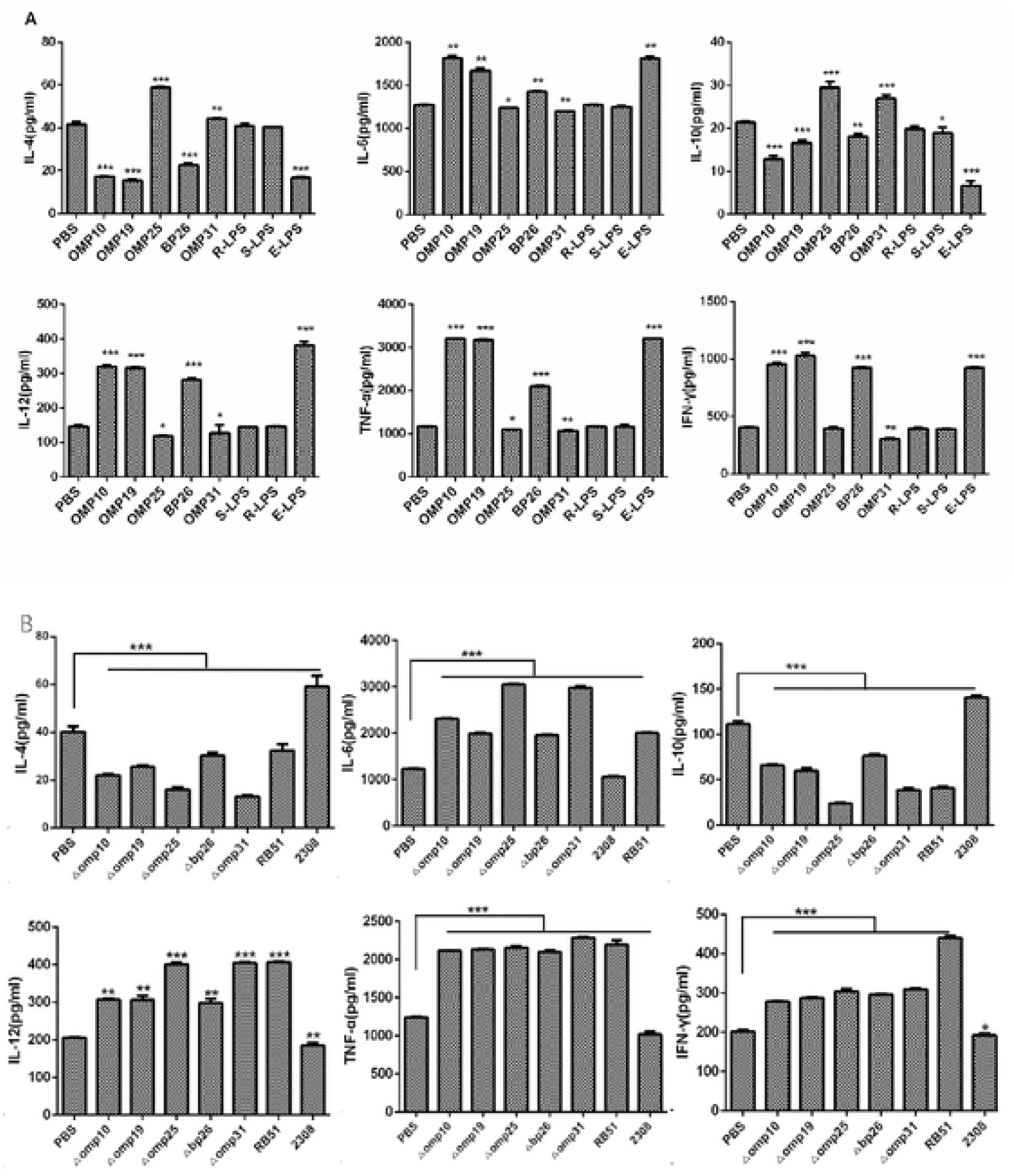
Cytokine secretion in supernatants of DCs infected with OMPs and *Brucella* mutants. DCs were cultured with purified OMPs (endotoxin removed), R-LPS, S-LPS, E-LPS (A), or *Brucella* mutants, *B. abortus* RB51, 2308 and PBS for 24 h (B), The culture supernatant was collected and the levels of secreted cytokines IL-4, IL-6, IL-10, IL-12, TNF-α and IFN-γ quantified by ELISA. Data shown are the means ± SEMs of three separate experiments. Significance was calculated using the t-test. * *P* < 0.05, ** *P* < 0.01 and *** *P* < 0.001.

### OMPs and *Brucella* mutants affect the transcription level of TLRs

Real-time PCR (RT-PCR) was used to determine TLRs mRNA expression. Only the expressions of TLR2 and TLR4 were enhanced significantly in OMPs (*P* < 0.05) (Fig. 3A), whereas the transcriptional levels of TLR2, TLR4 and TLR9 mRNA were enhanced significantly in *Brucella* mutants (*P* <0.05) (Fig. 3B). It has been suggested that TLR-mediated signaling pathways play a synergistic role in the maturation and antigen presentation of DCs.

**Fig. 3.**
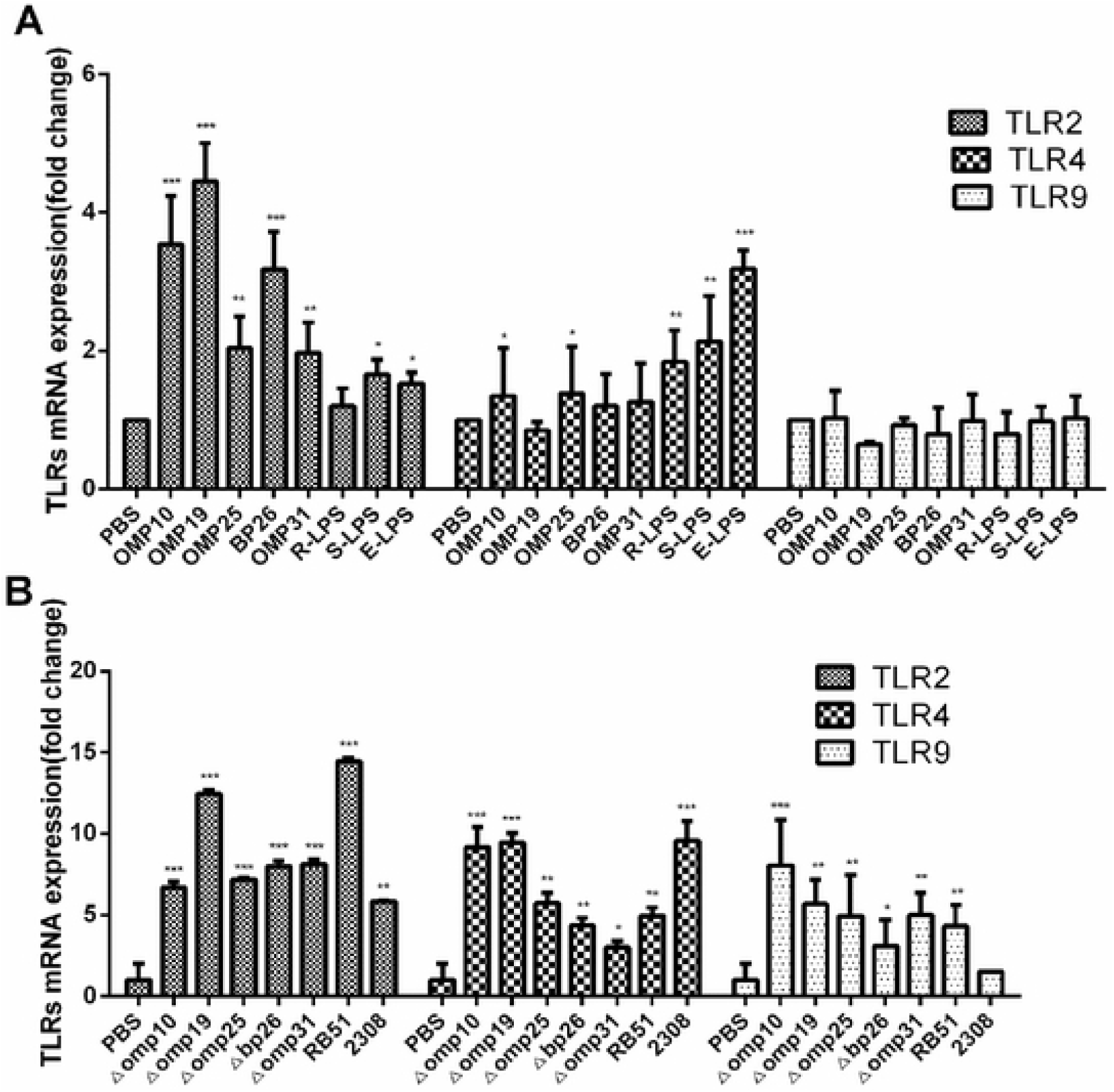
Effects of OMPs and *Brucella* mutants on transcription of TLRs mRNA in DCs. DCs were stimulated by OMPs, R-LPS, S-LPS, E-LPS (A), and *Brucella* mutants, *B. abortus* RB51 and 2308 (B) for 24 h. DCs were then collected, and total RNA was extracted for RT-PCR detection of TLR-2, TLR-4 and TLR-9 mRNA transcription levels. Statistical analysis was performed by the 2-ΔCT method. Experiments were performed three times in duplicate. * *P* < 0.05, ** *P* < 0.01, and *** *P* < 0.001.

### MTT assay

MTT colorimetric assay was used to measure T cell viability. Compared with the PBS group, DCs stimulated by OMP10, OMP19 and BP26 enhanced T lymphocyte proliferation more effectively, whereas DCs stimulated by OMP25 and OMP31 inhibited T cell proliferation (Fig. 4A). In addition, the proliferation efficiency of T cells in each group was the highest when the ratio of DCs: T cells was 1: 50. DC infection with *Brucella* mutants and B. abortus RB51 was more effective, and the proliferation efficiency was the most significant when the ratio of DCs: T cells was 1:25 (Fig. 4B). These results suggest that the proliferation efficiency of T cells stimulated by DCs is related to the antigen-presenting ability of DCs and the concentration of T cells.

**Fig. 4.**
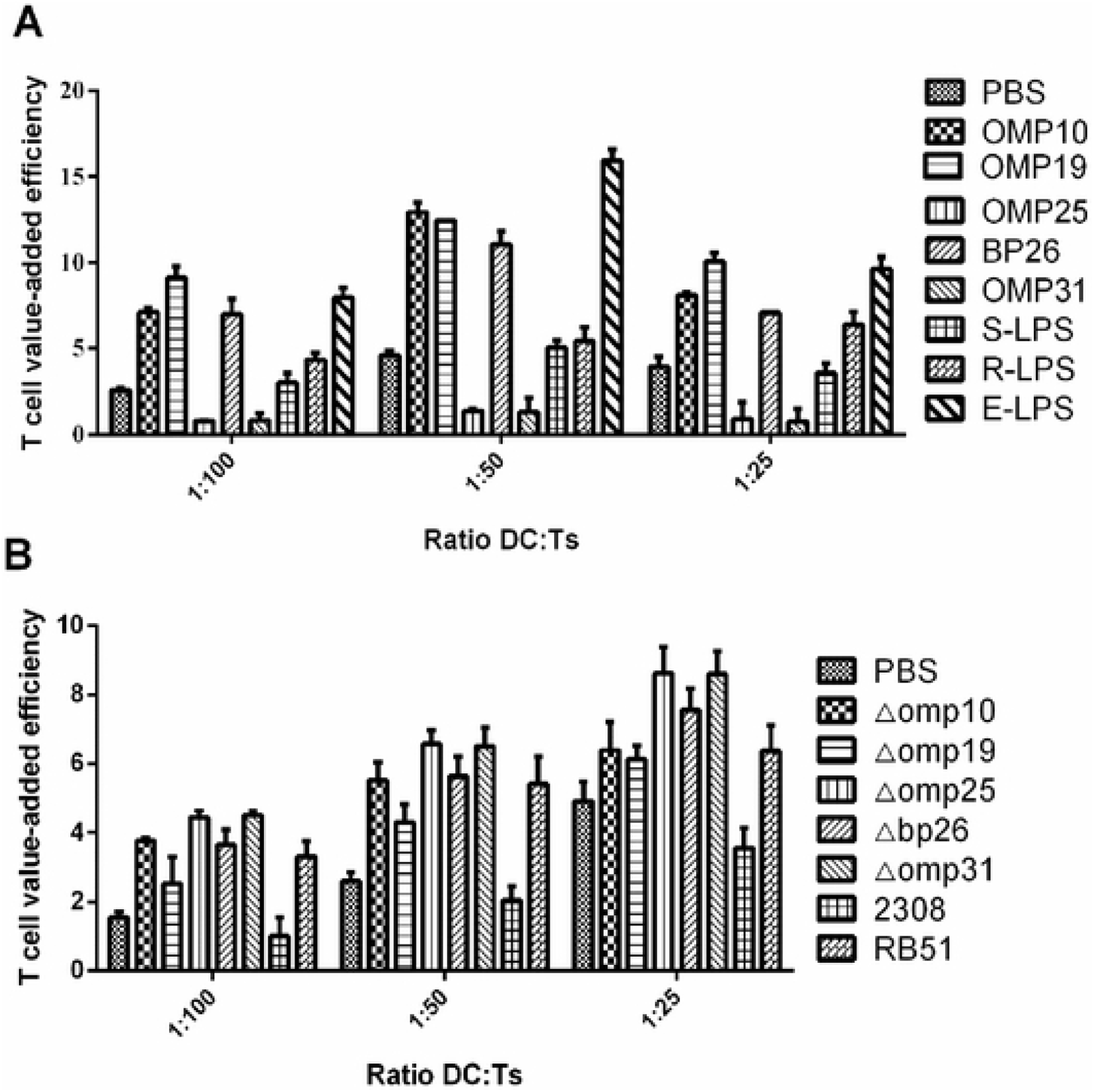
Effects of OMPs and *Brucella* mutant-stimulated DCs on T lymphocyte proliferation. DCs were stimulated with OMPs, R-LPS, S-LPS, E-LPS (A), or *Brucella* mutants, *B. abortus* RB51 and *B. abortus* 2308 (B) for 24 h, mixed with naive T cells and incubated at different concentrations (1:25, 1:50, 1:100) for 72 h. The proliferation efficiency of T lymphocytes in each group was examined. Each experiment was repeated at least three times.

### *Brucella* mutants can induce apoptosis of DCs

The effect of *Brucella* mutants on apoptosis of DCs were detected by flow cytometry. Compared with the PBS control group, the apoptosis level of DCs was dramatically increased in the *Brucella* mutants, *B. abortus* 2308 and *B. abortus* RB51 groups (*P* < 0.001) (Fig. 5A). The apoptosis rates were shown in the Fig. 5B. The results show that apoptosis is increased after stimulation of DCs with *Brucella* mutants and *B. abortus* RB51 and *B. abortus* 2308.

**Fig. 5.**
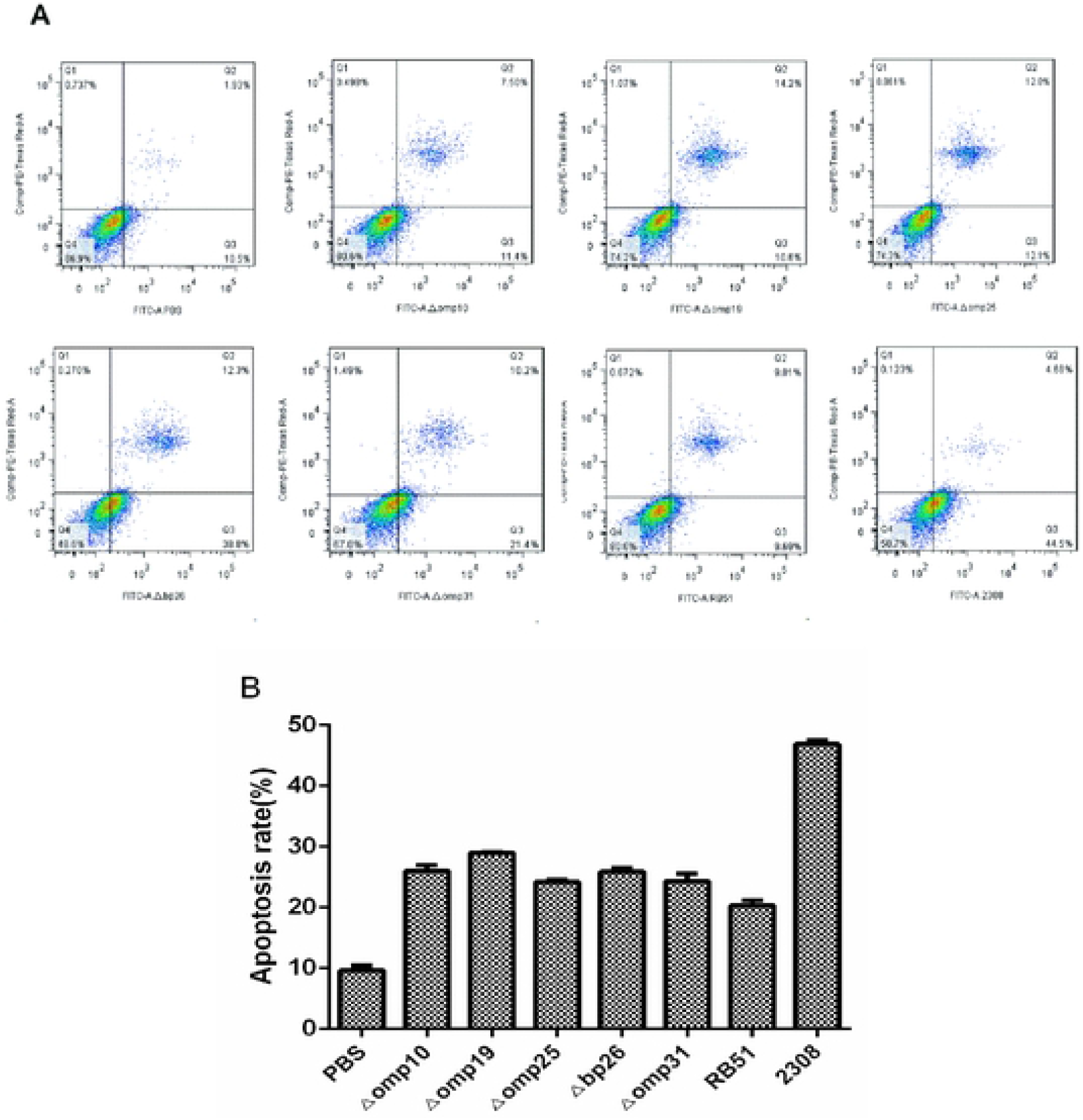
Apoptosis detection in DCs infected by *Brucella*. DCs were collected and stained by Annexin V/PI at 24 h post-infection and apoptosis rate was detected by flow cytometry. (A) Annexin V versus PI dot plots graphs. (B) GraphPad shows the result of one representative experiment of the three performed. Quantification of the percentage of Annexin V-positive cells. Bars express the mean ± SD of duplicates. Data shown are from a representative experiment of three independent experiments. * *P* < 0.05, ** *P* < 0.01, and *** *P* < 0.001.

### *Brucella* mutants decreased *Brucella* survival and replication in DCs

DCs were infected with *Brucella* and *Brucella* mutants and the survival capacity and replication capability of the Brucella in the DCs were determined. The DCs were infected with the seven strains at a MOI of 5, and the number of surviving bacteria was calculated. At 1 h post-infection, there was an overall decrease in the number of bacteria in DCs that infected with *Brucella* mutants (Δ*omp10*, Δ*omp19*, Δ*omp25*, Δ*omp31*) and B. abortus RB51 when compared with DCs that infected with *B. abortus* 2308. However, at 6 h post-infection, only the number of bacteria in DCs treated with mutants (Δ*omp19*, Δ*omp31*) and *B. abortus* RB51 had decreased when compared with *B. abortus* 2308. At 12 h post-infection bacteria number in DCs that received *Brucella* mutants (Δ*omp10*, Δ*omp19*, Δ*omp25*, Δ*omp31*) and *B. abortus* RB51 was significantly lower compared to DCs that infected with *B. abortus* 2308. At 24 h post-infection, there was an overall significant decrease in the number of bacteria in DCs that infected with *Brucella* mutants (Δ*omp19*, Δ*omp25*, Δ*omp31)* and *B. abortus* RB51 when compared with DCs that infected with *B. abortus* 2308. Finally, at 48 h post-infection, there was all significant decrease in the number of bacteria in DCs that infected with *Brucella* mutants (Δ*omp19*, Δ*omp26*) and *B. abortus* RB51 compared with DCs that infected with *B. abortus* 2308 (Fig. 6). These results suggest that the survival and replication ability of *Brucella* is decreased by *Brucella* mutants in DCs.

**Fig. 6.**
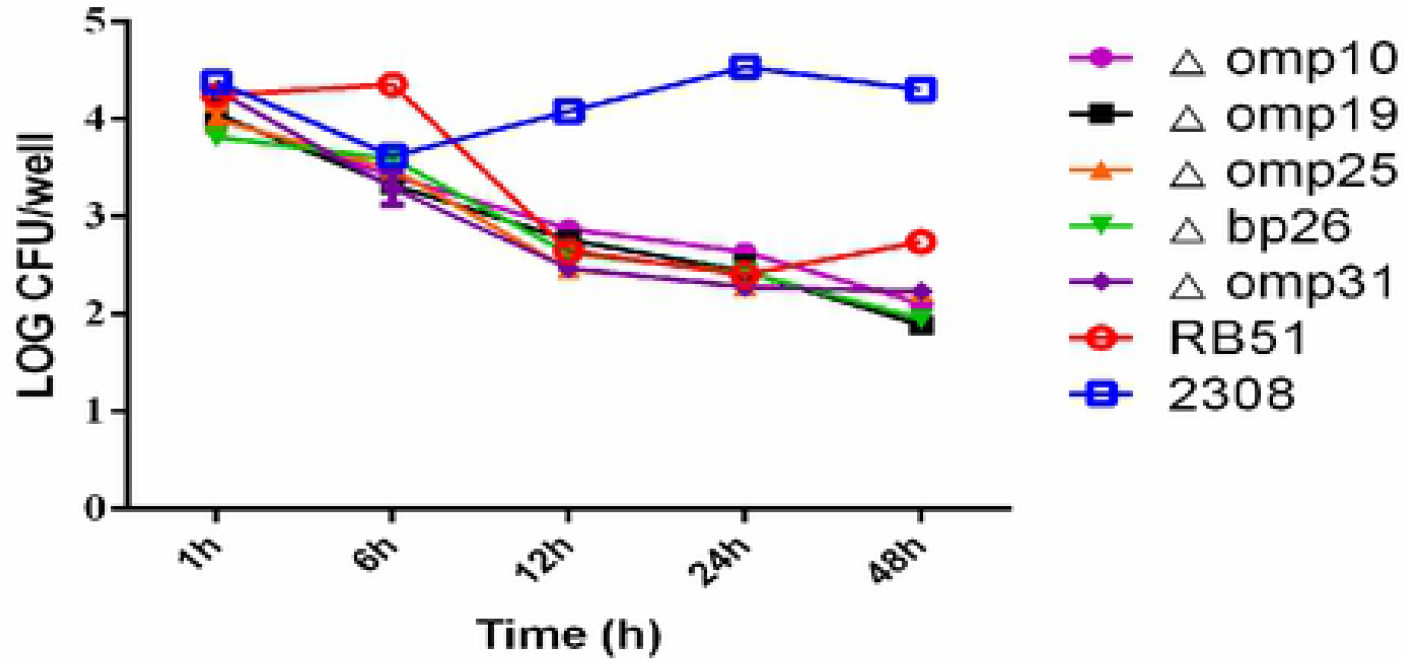
Survival capability and replication of *Brucella* in DCs. DCs were infected with *Brucella* mutants and *B. abortus*RB51, 2308 at a MOI of 5. At the indicated time points, DCs were lysed, and the bacterial count was quantified by plating serial dilutions on *Brucella* Agar plates. Three parallel controls were performed at each time point. CFU, colony forming units.

## Discussion

In this study, the effects of *Brucella* OMPs and *Brucella* mutants on the maturation and antigen presentation of DCs were evaluated. DCs were stimulated with *Brucella* OMPs and *Brucella* mutants and expression of the co-stimulatory molecule was assessed to investigate DC maturation and antigen presentation. The levels of cytokines in the supernatant were measured and the ability of DCs to shift T lymphocyte proliferation and differentiation was assessed, as well as mRNA expression levels of transcription factors TLR2, TLR4 and TLR9. DCs were stimulated by *Brucella* and *Brucella* mutants to evaluate the rate of apoptosis and intracellular survival in DCs. These results indicate that *Brucella* OMPs and *Brucella* mutants affect the maturation and antigen presentation of DCs.

DCs have the ability to express various pathogens recognition receptors (PRRs), which enable them to quickly sense stimulation by invasive pathogens and initiate an innate immune response [39]. However, DCs have established a special, adjustable phagocytic pathway, mainly by initiating antigen-specific immune responses to foreign antigens that destroy tissues and maintaining tolerance to self-antigens [40]. Over the past 10 years, DCs have been proven to be the key cell group determining the starting point of the specific immune response [41]. Moreover, when mature DCs present antigens to the initial T lymphocytes and stimulate their proliferation, they also induce the subsequent polarization of the adaptive immune response to the distribution of Th1 and Th2 cells by secreting different cytokines. For example, IL-4 stimulates initial T lymphocytes to differentiate into Th2 cells and regulates the humoral immune response [42,43]. IL-10 not only inhibits the expression of MHC-II molecules and co-stimulatory molecules, including CD40, CD80 and CD86, on the surfaces of macrophages and DCs but also inhibits the secretion of inflammatory cytokines (TNF-α, IL-12), thus inhibiting cellular antigen presentation [30]. Recent studies have found that *Brucella* regulates TNF-α secretion through an OMP25-dependent mechanism, inhibits DCs maturation and IL-12 secretion, and inhibits initial T cell activation and antigen presentation, thus inducing chronic infection [15]. It has been found that several *Brucella* proteins, including lipoprotein OMP19 [9], *Brucella* Lumazine synthase (BLS) [44], OMP16 [45], and BvrR [46] can induce DC maturation. These *Brucella* proteins may interact with DCs through an O antigen-independent pathway. In the current study, it was found that *Brucella* OMPs (OMP10, OMP19, BP26), *Brucella* mutants (Δ*omp10*, Δ*omp19*, Δ*omp25*, Δ*bp26*, Δ*omp31*), and *Brucella* RB51 induced the maturation and antigen presentation of DCs and improved activation of T lymphocyte proliferation, which is beneficial for recognizing antigens and eliminating pathogens, whereas *B. abortus 2308* reduced the DC maturation and antigen presentation to T cells by maintaining the balance between intracellular replication and inducing apoptosis, thus escaping the immune function of the host and establishing chronic infection.

In this study, the signal pathway mediated by toll-like receptors (TLRs) were found to be involved in the maturation and antigen presentation of DCs induced by *Brucella* OMPs and *Brucella* mutants. DC activation and maturation were initiated by recognition of pathogen-related molecular patterns (PAMP) and TLRs [15]. TLRs transduce signals through common junction molecules to stimulate DCs to secrete inflammatory factors [12]. Studies have shown that *Brucella* transduces signals through TLR2, TLR4 and MyD88 [47]. As clearance of *Brucella* by MyD88 is very important, it may be that TLR9 (also signal transduced through MyD88) plays a major role in the interaction between DCs and *Brucella* [15]. Zwerdling[48] et al also demonstrated that *Brucella* transduces signals through TLR2 and TLR4 in DCs. Oliveira [49] and Huang [50] et al demonstrated the crucial role of TLR9 in DC-mediated IL-12 secretion and *Brucella* clearance.

In macrophages, it has been reported that the B. abortus RB51 induces apoptosis, whereas the *B. abortus* 2308 inhibits apoptosis [51]. However, the current study found that B. abortus RB51 and *B. abortus* 2308, as well as the *Brucella* mutants induced apoptosis, and that *B. abortus* 2308 induced a significantly higher level of apoptosis. The *B. abortus* 2308 induced apoptosis at different status in macrophages and DCs, which may be related to the different immune types induced by the two host cells and the different interactions between *Brucella* and the two host cells [52]. Macrophages are full-time phagocytes that engulf and remove exogenous antigens and can expose antigens to the extracellular environment and facilitate host clearance of *Brucella* [53]. DCs are the main antigen-presenting cells, which process antigens and presents them to the surface of other cells in the immune system. Studies have demonstrated that apoptotic cells themselves can inhibit inflammation by sending inhibitor signals to DCs. Moreover, internalization of apoptotic cells don’t induce DCs maturation but maintains DCs in a semi-mature state [30]. Immature DCs migrate constitutively from the periphery to the lymphoid organs and present self-antigens to induce T cell tolerance. Studies have shown that DCs downregulate the expression of chemokine receptor CCR5 after uptake of apoptotic cells and up-regulation of CCR7 expression, thus transforming from inflammatory cytokines to lymphoid chemokines, resulting in DCs homing to draining lymph nodes [54]. After uptake of apoptotic cells, DCs not only maintain an immature phenotype but also secrete TGF-β1, which lead to the induction of regulatory T cell immunosuppression [30,55]. Therefore, DCs play an important role in the host immune response to pathogens and maintenance of tissue tolerance.

## Conclusions

In summary, these results show that *Brucella* OMPs (OMP10, OMP19, BP26, OMP25, OMP31) and *Brucella* mutants (Δ*omp10*, Δ*omp19*, Δ*omp25*, Δ*bp26*, Δ*omp31*) are involved in the regulation of DC maturation and antigen presentation. The above results can contribute to the rational design of vaccines against *Brucella* and a better understanding of the pathogenesis of *Brucella* and the protective immune response in the host. However, in vivo studies are needed on the specific mechanisms through which *Brucella* affects the maturation of DCs, the efficiency of antigen presentation and the stimulation of T cell proliferation.

## Acknowledgments

This study was supported by grants from National Natural Science Foundation of China (No. U1803236; 32002245), Major scientific and technological projects of the Corps (No. 2017AA003) and Shihezi University Youth Innovative Talent Cultivation Program (No. CXPY201909). We thank LetPub (www.letpub.com) for its linguistic assistance during the preparation of this manuscript. We are all grateful to the reviewers for their invaluable comments to improve this manuscript.

## Author Contributions

Conceptualization: BC, CC, ZW

Data curation: NY, ZT, MX

Formal analysis: MX, QZ, HZ, YW, YJ, TS

Funding acquisition: CC, ZW

Investigation: NY, ZT

Methodology: NY, BC, CC, ZW

Project administration: CC

Resources: CC, ZW

Software: NY, ZT

Supervision: CC, ZW

Validation: NY, BC, CC, ZW

Writing-original draft preparation: NY, ZT, ZW

Writing—review and editing: BC, CC, ZW

